# Distributed and gradual microstructure changes track the emergence of behavioural benefit from memory reactivation

**DOI:** 10.1101/2022.04.28.489844

**Authors:** Martyna Rakowska, Alberto Lazari, Mara Cercignani, Paulina Bagrowska, Heidi Johansen-Berg, Penelope A. Lewis

## Abstract

Memory traces develop gradually and link to neural plasticity. Memory reactivation during sleep is crucial for consolidation, but its precise impact on plasticity and contribution to long-term memory storage remains unclear. We used multimodal diffusion-weighted imaging to track the location and timescale of microstructural changes following Targeted Memory Reactivation (TMR) of a motor task. This showed continuous microstructure plasticity in precuneus across 10 days post-TMR, paralleling the gradual development of behavioural benefit. Both early (0 - 24 h post-TMR) and late (24 h - 10 days post-TMR) microstructural changes in striatum and sensorimotor cortex were associated with the emergence of behavioural effects of TMR at day 20. Furthermore, the baseline microstructural architecture of sensorimotor cortex predicted TMR susceptibility. These findings demonstrate that repeated reactivation of memory traces during sleep engenders microstructural plasticity which continues days after the stimulation night and is associated with the emergence of memory benefits at the behavioural level.

## 1 INTRODUCTION

The ability to change and adapt in response to internal or external stimuli by reorganising neural connections, structure, and function is a fundamental property of the brain ^1^. The remarkably plastic nature of the nervous system forms the basis of learning and memory ^2-4^. Learning-associated neural processes occurring during sleep have been receiving increasing attention in recent times. The active systems consolidation model suggests that newly encoded memories are reactivated during non-rapid eye movement (NREM) sleep, and that this enables their re-coding from a temporary store to a more permanent location ^5-7^. However, while the process of memory consolidation is gradual and occurs over long timescales ^8^, it is unclear whether reactivation of memories during sleep leads to plasticity in the neural substrate over time. Moreover, there is increasing evidence for distributed long-term cortical storage of memories ^9-11^, but it is unclear whether replay-driven consolidation of memories is associated with plasticity at different cortical sites. Likewise, how such plasticity could lead to long-term memory storage in humans has not been studied sufficiently ^12^.

Here we set out to track the location and timescale of microstructural changes underlying long-term effects of memory reactivation during sleep using Targeted Memory Reactivation (TMR). TMR involves associating learning items with sensory cues during wake and then covertly re-presenting them during sleep (e.g., ^13,14^). This is thought to trigger reactivation of the cue-associated memory representation which leads to a better recall of the cued items compared to those that were not cued during the night (i.e., uncued) ^15-19^. In recent years, TMR has become a valuable tool to study the mechanisms of sleep-dependent memory processes. It has allowed us to establish a causal link between memory reactivation and consolidation ^20,21^, and to identify brain regions that are functionally involved in such relationship ^13,18,22-24^. Our prior studies further demonstrated that the behavioural effects of TMR develop over time and can last up to three weeks ^19,24^. However, the physical brain changes driving the long-term functional and behavioural benefits of TMR remain poorly understood. We have recently shown that TMR can impact on grey matter volume within the task-related regions, thus shaping their functional response and the behavioural benefit of cueing in the long run ^24^. Yet, it is still unclear whether repeated reactivation of a memory trace can also modify tissue microstructure and what the time scale of such changes might be.

We used diffusion-weighted MRI (DW-MRI) to examine short- and long-term microstructural plasticity after TMR of a procedural memory task. DW-MRI is sensitive to the random motion of water molecules within tissue, thus providing indirect information about microstructure ^25^. It is a task-independent measure which, in contrast to functional MRI, allows probing of the microstructural substrate without being affected by task specific-activity. Importantly, the technique is also known to be sensitive to experience-driven plasticity within memory-related areas ^26-29^ and has been used to track the development of a neocortical engram, with the microstructural changes driving gains in behavioural performance ^30^. This makes DW-MRI an excellent technique for studying the gradual structural plasticity which is associated with memory formation over long timescales. The simplest approach to measure diffusion is diffusion tensor (DT) MRI, which assumes a Gaussian behaviour of water molecules within a single water compartment and allows the quantification of parameters such as mean diffusivity (MD) and fractional anisotropy (FA) ^31^. More complex models have been developed to account for the different behaviour of intra- and extra-cellular water, such as the Composite Hindered And Restricted Model of Diffusion (CHARMED ^32^). The relative sizes of the restricted (Fr, intracellular) and hindered compartments can then be estimated.

Our participants learned two motor sequences of a Serial Reaction Time Task (SRTT), each associated with a different set of auditory tones. Tones associated with one of the sequences were replayed to the participants during subsequent NREM sleep, successfully improving cued vs uncued sequence performance post-TMR (see ^24^ for behavioural effects). During both learning and the two re-test sessions (24 h and 10 days post-TMR), participants were placed in the scanner to acquire DW-MRI data, with the final re-test session taking place online, 20 days after TMR. MD has recently been shown to decrease in precuneus in response to a series of repeated learning-retrieval epochs during wake ^30^, which could be regarded of as a proxy of memory reactivation during sleep ^33^. We thus hypothesised that TMR during sleep would also lead to rapid MD-driven plasticity within precuneus, thereby supporting the functional activation of this structure in association with TMR that we observed in our previous work ^24^. However, we expected the motor-related regions to undergo long-term microstructural changes, thereby reflecting their slowly evolving reorganisation ^34-38^, as well as long-term functional engagement and volumetric increase in response to TMR ^24^. In the current study, we also used Restricted Water Fraction (Fr), as modelled by the CHARMED framework ^32^. Fr is thought to be more sensitive to microstructural changes than MD, especially in long-term assessment ^28^. We thus combined MD with Fr in a multimodal analysis protocol to uncover common microstructural patterns across the two MRI markers and their relationship with TMR benefits across time. Finally, despite the importance of tissue microstructure for memory formation ^30^, it is unclear whether baseline tissue microstructure is predictive of memory encoding capacity. To clarify this, we tested whether baseline brain characteristics can determine TMR susceptibility, thus adding to our current understanding of the factors that influence the effectiveness of TMR ^39^.

## 2. RESULTS

Post-sleep SRTT re-test sessions took place 24.67 hours (SD: 0.70) (Session 2, S2), 10.48 days (SD: 0.92) (Session 3, S3), and 20.08 days (SD: 0.97) (Session 4, S4) after Session 1 (S1). As we have reported previously ^24^, analysis of behavioural data showed a main effect of TMR on the SRTT reaction time performance (p = 0.001), with the difference between the cued and uncued sequence strongest at S4, i.e., 20 days post-TMR (p_adj_ = 0.004, Fig.1a). Furthermore, there was a main effect of the amount of time post-TMR on cueing benefit (p = 0.046, Fig.1a), suggesting that the effects of our manipulation develop over time before they emerge at S4 ^24^.

**Fig. 1.**
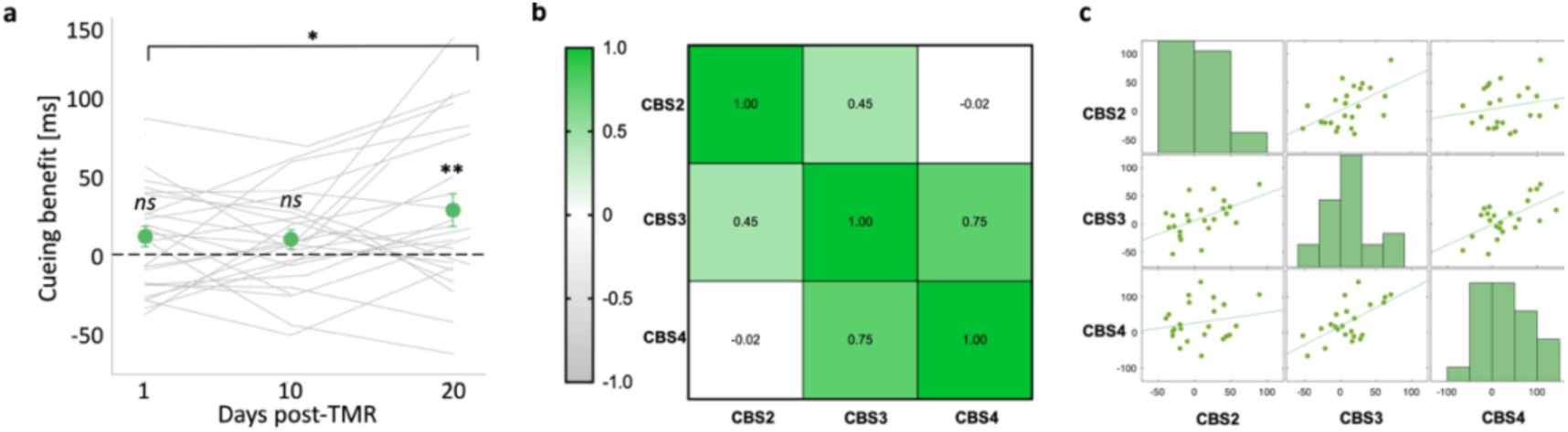
Behavioural effects of TMR. **(a)** Difference between the late sequence specific skill of the cued and uncued sequence (i.e., the cueing benefit) plotted against the number of days post-TMR. Green dots represent mean ± SEM calculated for S2 (1 day post-TMR), S3 (10-14 days post-TMR) and S4 (16-21 days post-TMR). Grey lines represent cueing benefit for each subject. A linear mixed effects analysis showed a main effect of time on cueing benefit, which itself was significant at S4. * p < 0.05;** p_adj_ = 0.004. **(b)** Heatmap of Pearson’s correlation coefficient matrix showing the relationships between cueing benefit (CB) at different sessions. **(c)** Scatterplots showing the same correlations as in (b). CBS2, CBS3, CBS4: cueing benefit at S2, S3, S4; S1-4: Session 1-4; For (a): n = 30 for S2, n = 25 for S3; n = 24 for S4. For (b-c): n = 23.

### 2.1 TMR-RELATED PLASTICITY

We sought to examine whether repeated reactivation of a motor memory trace could engender microstructural plasticity in the brain. We tested whether behavioural cueing benefit at different sessions is associated with changes in brain microstructure within predefined regions of interest (ROIs) over set periods of time. Thus, individual modality maps collected at different sessions were subtracted from each other, yielding measures of early (S1-S2) and late (S2-S3) microstructural plasticity. The resultant difference maps were used as inputs in the multimodal analysis, set out to uncover common trends in MD and Fr change measures. Cueing benefit at S2, S3 and S4 were entered as regressors, and separate contrasts were run for cueing benefit at each session. Given the correlations between cueing benefits at different sessions (Fig.1b-c), sessions of no interest in any given contrast were covaried out to test if the effects are specific to the cueing benefit tested.

We hypothesised that TMR during sleep would lead to rapid plasticity within precuneus. Consistent with this, the analysis revealed a positive relationship between early plasticity in left precuneus (−6, - 60, 18) and cueing benefit at S3 when controlling for the behavioural effects at S2 and S4 (p = 0.041; Fig.2a-c, Table S1A, Fig.S3a), such that greater cueing benefit was associated with greater reductions in MD and greater increases in Fr. In addition, late plasticity in bilateral precuneus (4, -58, 16) was associated with cueing benefit at S4 when controlling for the behavioural effects at S2 and S3 (p = 0.027; Fig.2d-f, Table S1B, Fig.S3b), again reflecting greater cueing benefit being associated with greater reductions in MD and greater increases in Fr. These results demonstrate that TMR can impact on precuneus microstructure both early and late in the consolidation process, thus supporting the development of behavioural effects of TMR over time.

**Fig. 2.**
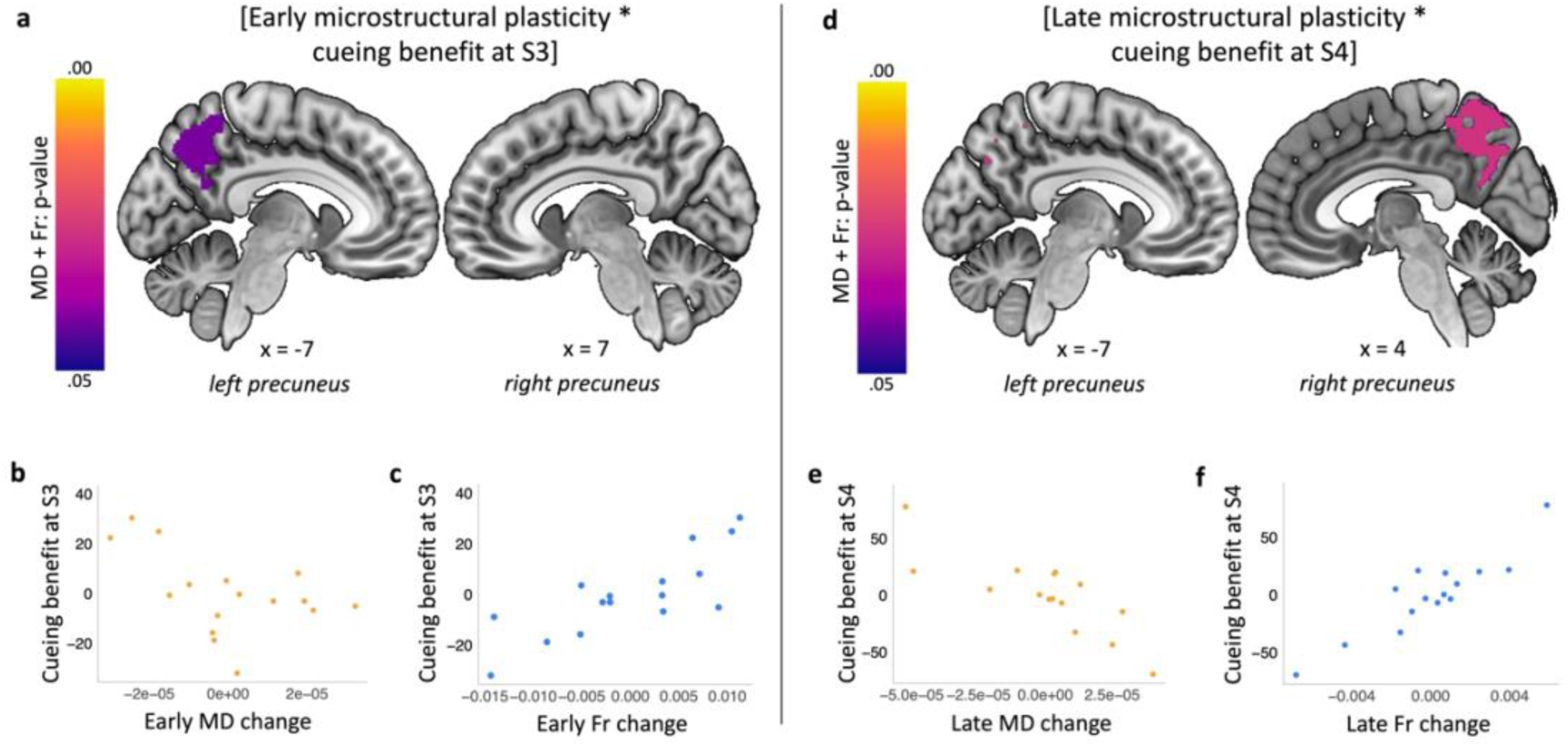
Multimodal plasticity in precuneus is associated with long-term cueing benefits. **(a)** Early (from S1 to S2) microstructural plasticity in left precuneus is associated with cueing benefit at S3. **(d)** Late (from S2 to S3) microstructural plasticity in bilateral precuneus is associated with cueing benefit at S4. Purple-yellow colour bars indicate p-values for the results thresholded at a significance level of p_FWE_ < 0.05, corrected for multiple voxel-wise comparisons within pre-defined ROI for bilateral precuneus. Results are overlaid on a Montreal Neurological Institute (MNI) brain. (**b, c, e, f**) Mean MD (b, e) and Fr (c, f) change values extracted from multimodal clusters significant at pFWE < 0.05 shown in (a) and (d). Scatterplots are presented for visualisation purposes only and should not be used for statistical inference. Each data point represents a single participant, axes represent residual values after correcting for age, sex, PSQI score, baseline reaction time, baseline learning capabilities on the SRTT, cueing benefit at S2 and either cueing benefit at S4 (b-c) or at S3 (e-f). S1-4: MD: Mean Diffusivity; Fr: Restricted Water Fraction; PSQI: Pittsburgh Sleep Quality Index; S1-S4: Session 1-4; n = 16 for (a-c), n = 15 for (d-f).

**Fig. 3.**
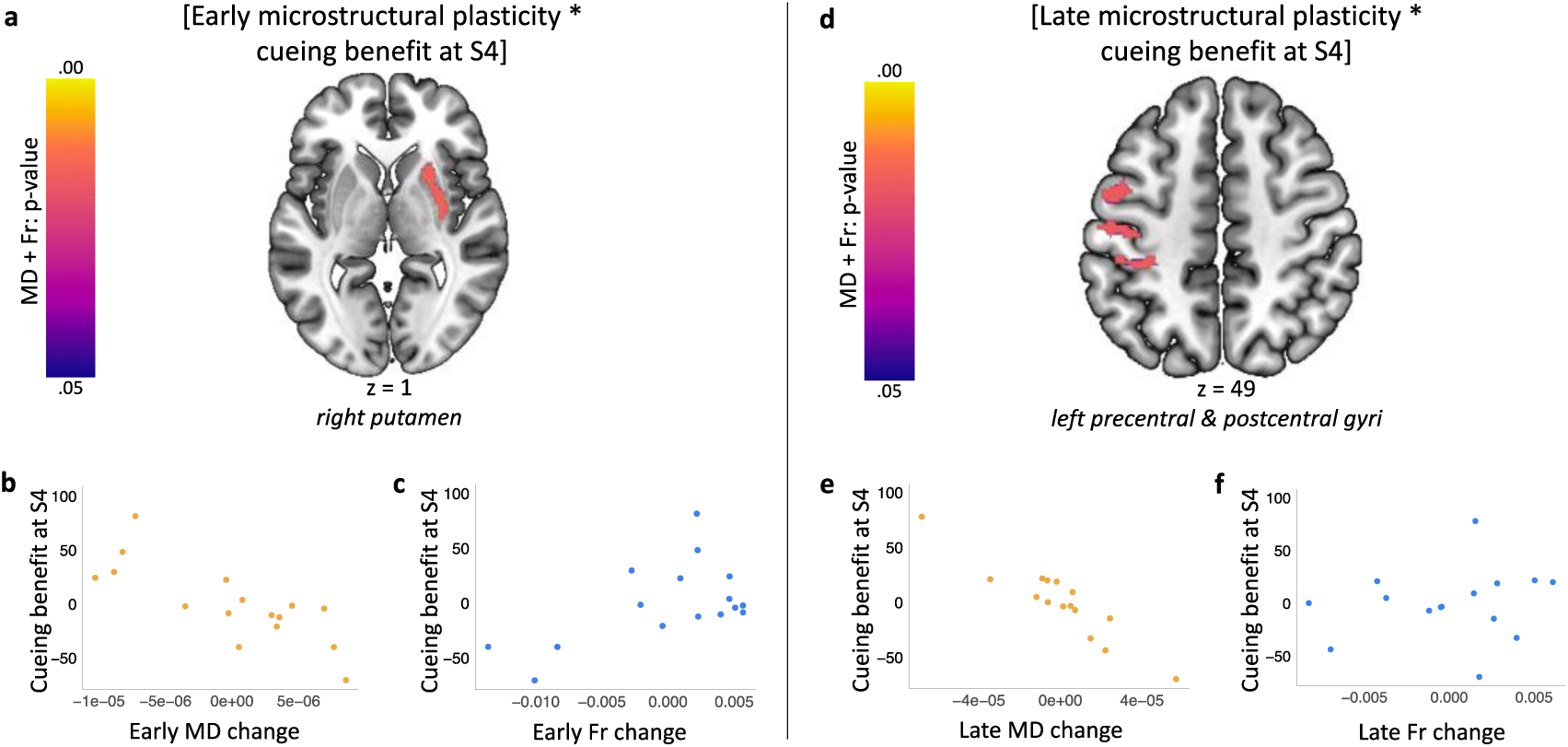
Multimodal plasticity within task-related structures is associated with long-term cueing benefits. **(a)** Early (from S1 to S2) microstructural plasticity in right putamen correlates with cueing benefit at S4. **(d)** Late (from S2 to S3) microstructural plasticity in left sensorimotor cortex correlates with cueing benefit at S4. Purple-yellow colour bars indicate p-values for the results thresholded at a significance level of p_FWE_ < 0.05, corrected for multiple voxel-wise comparisons within pre-defined bilateral ROI for putamen (a) and sensorimotor cortex (d). Results are overlaid on a Montreal Neurological Institute (MNI) brain. (**b, c, e, f**) Mean MD (b, e) and Fr (c, f) change extracted from multimodal clusters significant at p_FWE_ < 0.05 shown in (a) and (d). Scatterplots are presented for visualisation purposes only and should not be used for statistical inference. Each data point represents a single participant, axes represent residual values after correcting for age, sex, PSQI score, baseline reaction time, baseline learning capabilities on the SRTT, and cueing benefit at S2 and S3. MD: Mean Diffusivity; Fr: Restricted Water Fraction; PSQI: Pittsburgh Sleep Quality Index; S1-4: Session 1-4; n = 16 for (a-c), n = 15 for (d-f).

We further hypothesised that task-related sensorimotor regions would undergo longer-term microstructural changes. Our multimodal voxel-wise ROI analyses revealed that both early plasticity in right putamen (34, -6, -8) and late plasticity in left sensorimotor cortex (−60, -18, 14) were associated with cueing benefit at S4 (putamen: p = 0.016; Fig.2a-c, Table S1C, Fig.S3c; sensorimotor cortex: p = 0.018; Fig.2d-f, Table S1D, Fig.S3d). To confirm that the results were specific to S4, we also tested the relationship between microstructural plasticity and cueing benefit at S2 and S3. In line with our expectations, there was no relationship between motor network plasticity and cueing benefit at S2 or S3 (p > 0.05).

Together, these results provide evidence for gradual plasticity in the microstructure of precuneus, striatum and sensorimotor cortex that underpins long-term behavioural effects of TMR. Interestingly, both the early and late plasticity results for motor ROIs seem to be specifically related to S4, the time point at which the behavioural impact of TMR emerged.

### 2.2 INDIVIDUAL DIFFERENCES IN BASELINE MICROSTRUCTURE

A wide variety of factors are known to influence TMR’s success ^39^. We were interested to determine if inter-individual variability in brain microstructure could confer susceptibility to the manipulation. For this, we were not interested in the TMR effect at any particular session, but rather in the common variance of the cueing benefit shared across the post-stimulation sessions (Fig.1b-c). Thus, we performed a Principal Component Analysis (PCA) on the cueing benefit at S2, S3 and S4 in order to obtain a single measure which we will call ‘TMR susceptibility’. We used baseline (S1) maps of MD and Fr as inputs, with TMR susceptibility entered as a regressor. This showed a relationship between baseline microstructure in right precentral and postcentral gyrus (58, -6, 20) and TMR susceptibility (p = 0.041; Fig.4a-c, Table S2; Fig.S4), such that individuals with greater response to TMR had less MD and more Fr at baseline. This finding suggests that the individual variation in sensorimotor microstructure could predict susceptibility to TMR of procedural memory. Importantly, this finding is independent of participants’ demographics (sex, age), general sleep patterns (as measured by the Pittsburgh Sleep Quality Index (PSQI) score), time spent in stage 2 of NREM sleep (N2), baseline reaction time and learning capabilities on the SRTT, all of which were controlled for in the analysis.

**Fig. 4.**
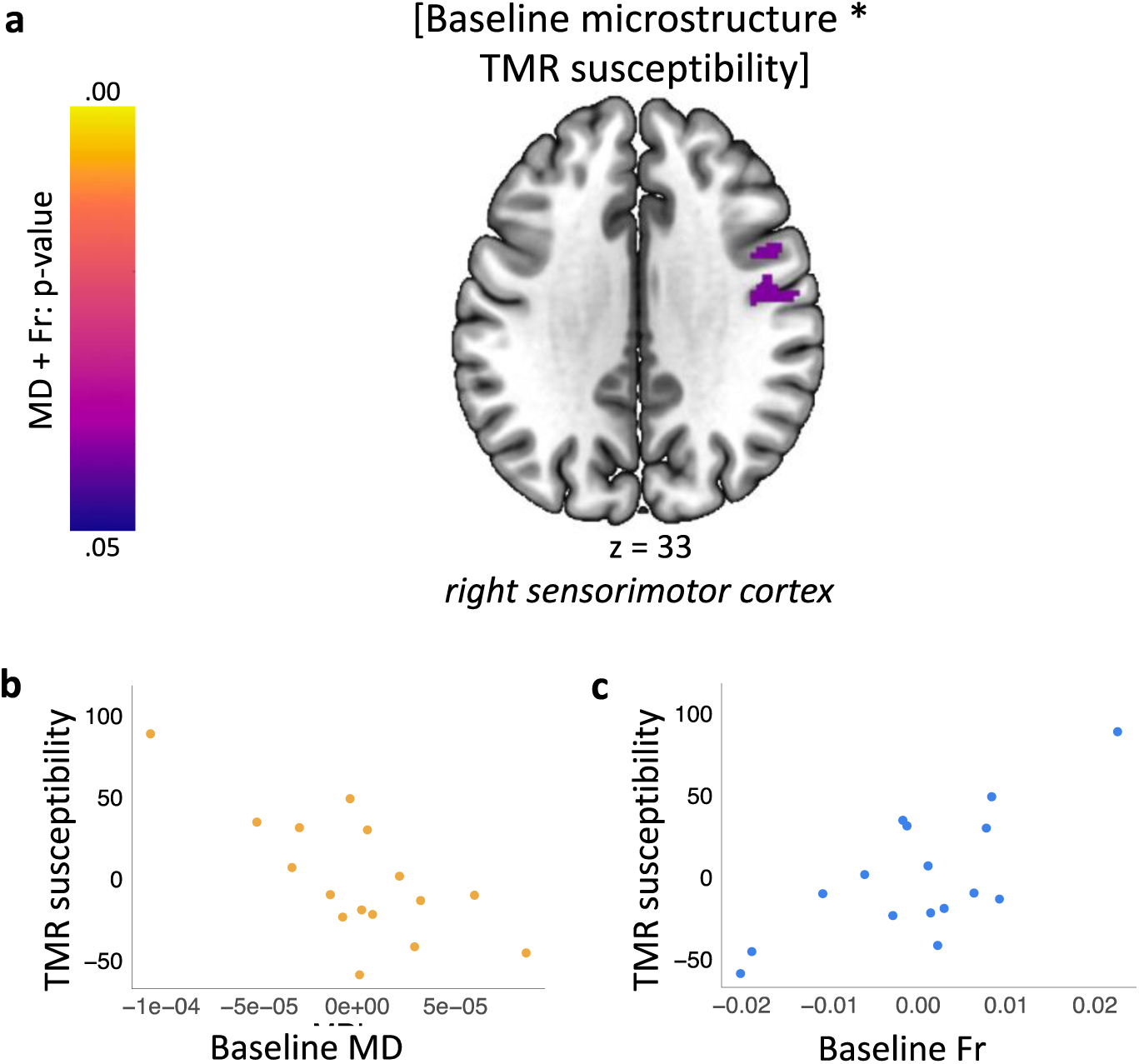
TMR susceptibility is associated with baseline sensorimotor microstructure. **(a)** Results of the multimodal analysis testing the relationship between TMR susceptibility and baseline microstructure. Colour bars indicate p-values for the results thresholded at a significance level of p_FWE_ < 0.05 (red), corrected for multiple voxel-wise comparisons within pre-defined ROI for bilateral sensorimotor cortex. Results are overlaid on a Montreal Neurological Institute (MNI) brain. **(b-c)** Mean baseline MD (b) and Fr (c) extracted from the clusters significant at p_FWE_ < 0.05 shown in (a). Scatterplots are presented for visualisation purposes only and should not be used for statistical inference. Each data point represents a single participant, axes represent residual values after correcting for age, sex, PSQI score, percentage of time spent in N2, baseline reaction time and baseline learning capabilities on the SRTT. MD: Mean Diffusivity; Fr: Restricted Water Fraction; PSQI: Pittsburgh Sleep Quality Index; S1-4: Session 1-4; n = 16.

## 3 DISCUSSION

We set out to investigate the relationship between plasticity in tissue microstructure and beneficial effects of memory reactivation during sleep. To this end, we combined TMR with DW-MRI to test whether baseline micro-architecture of the brain can be used to determine one’s susceptibility to TMR, and whether TMR can impact on brain microstructure, thus giving rise to the behavioural effects observed. First, we find that long-term cueing benefit is associated with gradual microstructural plasticity within memory and task-related regions. Specifically, our data suggests that precuneus undergoes microstructural plasticity throughout the consolidation process, supporting TMR-related benefits 10 and 20 days after the manipulation, respectively. In addition to precuneus, early microstructural changes in striatum and late microstructural changes in sensorimotor cortex also relate to the beneficial effects of cueing 20 days post-TMR. Second, we demonstrate that individual differences in baseline sensorimotor microstructure predict behavioural TMR susceptibility, i.e., the variance shared between cueing benefit at the combined post-TMR sessions.

### 3.1 TMR-INDUCED PLASTICITY IN PRECUNEUS

We show a relationship between both early (0 - 24 h post-TMR) and late (24 h - 10 days post-TMR) precuneus plasticity and cueing benefit 10 days and 20 days post-TMR, respectively. These results suggest that repeated reactivation of a memory trace during sleep could engender gradual microstructural changes within precuneus that are associated with the gradual emergence of behavioural benefits from the manipulation. We have previously shown that TMR of a procedural memory also engages precuneus functionally, but this occurs relatively early in the consolidation process ^19^. Given the well-described role of precuneus in memory retrieval ^40,41^, our previous results could reflect the difference in recall strength of cued and uncued sequences during their execution. Indeed, TMR and memory reactivation per se share a lot of parallels with memory retrieval. However, the long-term time scale of our current results as well as the microstructural changes that we report suggest that the role of precuneus may extend beyond retrieval only. We believe that precuneus could act as an ‘episodic buffer’ ^42^, building up representations of the retrieved information before they are transferred to a more permanent store. Indeed, precuneus has already been shown to undergo rapid, learning-dependent microstructural plasticity, indicative of memory engram development within this region ^30^. The structure is known to harbour behaviourally relevant memory representations ^43,44^ and its function is traditionally associated with motor learning ^45,46^. Our current results expand the existing literature by demonstrating that memory reactivation during sleep mediates microstructural plasticity which continues days after a single night of cueing within precuneus, perhaps facilitating the development of a stable and long-lasting memory trace in this structure. This, in turn, gives rise to the emergence of behavioural benefits of reactivation, reflected in the difference between the cued and uncued sequence that we observe 20 days post-TMR. Thus, we argue that (targeted) memory reactivation during sleep has a powerful impact on both memory processing and brain plasticity, and that its effects extend beyond the initial night of sleep.

### 3.2 NEUROPLASTICITY AND MICROSTRUCTURE OF THE MOTOR SYSTEM

Our findings support the suggestion that the long-term cueing benefit of TMR is mediated by early plasticity in putamen and late plasticity in precentral and postcentral gyri. We show that changes within these structures are specifically related to the behavioural effects 20 days post-TMR, suggesting that such microstructural plasticity in the motor system may give rise to the emergence of behavioural TMR effects. Both striatum and sensorimotor structures are thought to be critical for long-term storage of motor sequences ^47^. We build on this literature by arguing that memory reactivation during sleep may engender microplasticity within these regions, and thus stabilise memory traces harboured by the cortico-striatal system, shaping the sleep-dependent procedural benefits. Memory reactivation has been observed in ventral striatum immediately after learning ^48^ and this could drive the early microstructural plasticity in the adjacent regions, including putamen. In turn, the late microstructural plasticity that we observe in primary motor and somatosensory cortices likely reflects their slowly evolving reorganisation ^34-38^. This late plasticity could also underpin the TMR-related functional engagement and grey matter volume increase of the sensorimotor cortex which we observed previously ^24^. Interestingly, a recent rodent study found that cortico-striatal functional coupling increases during offline periods of rest and is required for long-term skill learning ^49^. Furthermore, this coupling seems to be mediated by NREM sleep spindles ^49^, which are known to be involved in motor learning ^50^. One intriguing possibility is that neuronal ensembles within sensorimotor cortex and striatum undergo simultaneous replay during post-learning sleep and this leads to their functional coupling. Simultaneous activity in primary motor cortex and dorsal striatum has already been recorded during motor learning ^51^. Our results raise the hypothesis that memory reactivation could also co-occur in these regions during sleep and thereby drive the synaptic plasticity within the underlying substrate. If this is correct, it might suggest that such co-replay could underpin the microstructural changes and the subsequent behavioural benefits observed in our dataset.

We further show that individual differences in the microstructural architecture of sensorimotor cortex measured at baseline are associated with TMR susceptibility. Thus, our results combine to suggest that the microstructure of sensorimotor cortex can both change in response to cueing motor memory reactivation and confer susceptibility to the stimulation. The success of TMR may therefore depend on the inter-individual variation in the microstructure of precentral and postcentral gyri. That is, the intrinsic micro-architecture of these task-related regions may either control memory encoding capacity, impact on the response to the manipulation or determine the effectiveness of the reactivation process itself. This finding adds to the existing literature on the factors modulating TMR’s success ^39,52-54^, perhaps explaining some of the discrepancies in the TMR literature.

### 3.3 BIOLOGICAL INTERPRETATION OF THE DW-MRI FINDINGS

The results of the current study demonstrate that DW-MRI can provide a valuable tool to investigate behaviourally relevant changes in brain microstructure. Furthermore, the multimodal approach that we adopted here revealed a common pattern across two diffusion markers: MD and Fr. This not only makes our findings more robust but also provides insights into the biological changes that could underpin sleep-dependent memory consolidation. Biological interpretation of diffusion measures is not straightforward ^2^, but combining multiple modalities increases the chances of picking up features that are shared by the two markers. In case of MD and Fr, water diffusion within the restricted (intracellular) volume fraction seems to be a common feature that both markers are sensitive to. Thus, the microstructural changes associated with memory reactivation could involve remodelling of the cylindrical tissue compartments, such as neural and glial processes ^28^. Indeed, rapid structural modifications after learning have been reported in astrocytic processes ^55,56^ and dendritic spines ^57^. In fact, both NREM ^58^ and REM sleep ^59^ have been implicated in dendritic spine plasticity within hours after learning, while disrupting memory reactivation during sleep impaired post-training spine formation ^58^. By the same token, repeated reactivation through TMR during sleep could have boosted similar forms of plasticity in the cylindrical compartments of precuneus, striatum and sensorimotor cortex in our dataset, thus giving rise to the observed changes in the DW-MRI metrics. However, swelling of cells (particularly astrocytes ^60^), and thus a shift in the ratio of extra-to intra-cellular space ^55,61,62^ could also alter the diffusion properties of the tissue and, consequently, the MD and Fr values. Indeed, synaptogenesis ^35^ and astrocytic hypertrophy ^63^ are detectable only after 7-10 days of training, thus matching the time scale of our late microstructural plasticity. Nevertheless, the cellular processes driving MD and Fr changes are generally difficult to identify and histological approaches would be needed to confirm the biological interpretation of our results.

In a broader context, DW-MRI allowed us to study the dynamic and distributed nature of the TMR-related changes. Our results demonstrate that cued memory reactivation can modify tissue microstructure. Notably, we show the resulting plasticity encompasses several cortical areas, continues days after the stimulation nights and supports long-term consolidation of memories that are reactivated during sleep. Thus, we extend the existing literature by providing direct evidence for reactivation-mediated redistribution of memory traces across the brain ^6,64^. Given the long-term character of the detected changes, we speculate that cueing memory reactivation must have either primed the relevant synapses for later processing ^64^ or biased plasticity-related protein capture towards the targeted memory traces ^65^. The consequential bias in synaptic plasticity would explain why a single night of cueing was capable of modifying tissue microstructure up to 10 days later. We further identify precuneus and motor structures as important neocortical memory hubs for long-term retention of procedural memories. The microstructural changes in precuneus, striatum and sensorimotor cortex were specifically related to the behavioural effects 20 days post-TMR, which was the only time point where we find group level evidence for the difference between the cued and uncued sequence ^24^. This suggests that the microstructural plasticity continuously parallels the gradual development of behavioural cueing benefit, eventually contributing to the emergence of behavioural effects of TMR.

## 4 CONCLUSION

We show that gradual microstructural changes, distributed across several cortical areas, track the emergence of behavioural benefits stemming from memory reactivation. Specifically, we find that microstructural plasticity occurs in precuneus and the motor system over different periods of time and is associated with long-term benefits of procedural memory TMR. These findings support the long-lived belief that stable memory traces develop gradually and reorganise the underlying tissue ^8^. Our results were specific to the cueing benefit 20 days post-manipulation, suggesting that the microstructural changes observed are specifically linked to the behavioural gain from memory reactivation. They also demonstrate that DW-MRI can be used to detect behaviourally relevant processes that underpin sleep-dependent memory consolidation. Finally, we shed new light on the factors that influence TMR’s effectiveness by demonstrating that individual variation in the microstructure of the task-related regions can be used to predict one’s susceptibility to the manipulation.

## 5 MATERIALS AND METHODS

### 5.1 PARTICIPANTS

The same sample of 33 healthy volunteers that we reported previously ^24^ signed a written informed consent to take part in the study, which had been approved by the Ethics Committee of the School of Psychology at Cardiff University. All participants reported being right-handed, sleeping approximately 8 h per night, having normal or corrected to normal vision, no hearing impairment, and no prior knowledge of the tasks performed upon the start of the study. Regular nappers, smokers, subjects who had travelled across more than two time-zones or engaged in any regular night work during one month prior to the experiment were not recruited. Further criteria for exclusion included recent stressful life event(s), regular use of any medication or substance affecting sleep, prior history of drug/alcohol abuse, and neurological, psychological, or sleep disorders. Additionally, participants were asked to abstain from napping, extreme physical exercise, caffeine, alcohol, and other psychologically active food from 24 h prior to each experimental session. We also excluded participants with more than three years of musical training in the past five years due to a probable link between musical abilities and procedural learning ^66,67^. Participants were screened by a qualified radiographer from Cardiff University to assess their suitability for MRI and signed an MRI screening form prior to each scan.

Four participants had to be excluded from all analyses due to: technical issues (n = 1), voluntary withdrawal (n = 1), interrupted electroencephalography (EEG) recording during the night (n = 1), and low score on the handedness questionnaire (indicating mixed use of both hands), combined with positive slope of learning curve during the first session (indicating lack of sequence learning before sleep) (n = 1). Six additional participants had to be removed from the DW-MRI analyses due to failure of the posterior (n = 2) or anterior (n = 4) part of the radiofrequency coil. Hence, 23 participants remained in the final dataset (12 females, age range: 18 -23 years, mean ±SD: 20.42 ± 1.56; 11 males, age range: 19 - 22 years, mean ±SD: 20.18 ± 0.98). Due to COVID-19 outbreak, four participants were unable to complete the study, missing either one (n = 1) or two (n = 5) sessions. One additional participant could not physically attend S3; they performed the SRTT online, but their MRI data could not be collected and therefore the sample size for the MRI analyses of S3 had to be decreased by one. Finally, Fr maps collected from three additional participants failed a visual quality check after pre-processing and were thus excluded from the Fr analysis (n = 3). Hence, the final sample size for MD analysis was n = 23 for S1, n = 23 for S2 and n = 19 for S3, and the sample size for Fr analysis was n = 20 for S1, n = 20 for S2 and n = 16 for S3. A flowchart of participants included and excluded from the different analyses is presented in Fig.S1.

### 5.2 STUDY DESIGN

The study consisted of four sessions (Fig.5a), all scheduled for the same time in the evening (∼ 8 pm). Upon arrival for the first session (S1), participants completed Pittsburgh Sleep Quality Index (PSQI) ^68^ to examine their sleep quality over the past month. S1 consisted of a motor sequence learning task (the SRTT), MRI data acquisition and overnight stay in the lab. The SRTT learning session was split in half, such that the first half of the SRTT blocks (24 sequence blocks) was performed in a 0T Siemens ‘mock’ scanner (i.e., an environment that looked exactly like an MRI scanner, but with no magnetic field) and the other half (24 sequence blocks + 4 random blocks) in a 3T Siemens MRI scanner, immediately after T1-weighted (T1w) structural data acquisition. This was followed by DW-MRI (see section 2.*6 MRI data acquisition*). Participants were then asked to prepare themselves for bed and were fitted with an EEG cap. While in stable stage 2 (N2) or 3 (N3) of NREM sleep, the TMR protocol was initiated (see section *2.5 TMR during NREM sleep*). Briefly, to trigger reactivation of the associated SRTT memories, tones associated with one of the SRTT sequences were replayed to the participants through speakers (Harman/Kardon HK206, Harman/Kardon, Woodbury, NY, USA). After, on average, 8.81 ±0.82 hours in bed participants were woken up and had the EEG cap removed before leaving the lab.

**Fig. 5.**
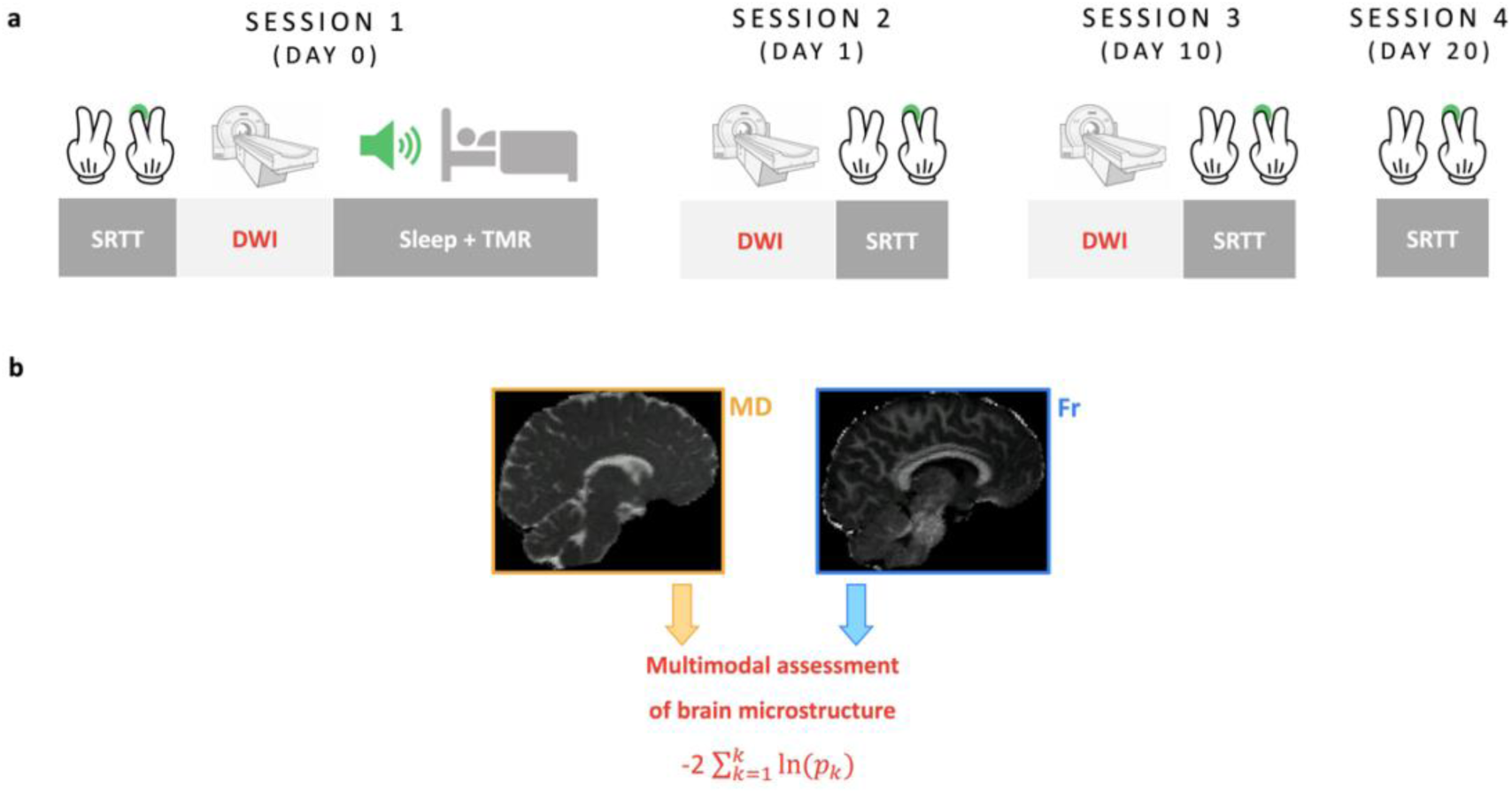
Experimental methods. **(a)** Study design. The study consisted of four sessions, each requiring participants to complete the SRTT. During S1, SRTT was followed by DW-MRI acquisition. During S2 (24 h post-TMR) and S3 (10 days post-TMR), the order was flipped, with the SRTT following MRI data collection. Structural T1w data was acquired at the beginning of each scan (S1-S3). S1 also involved EEG recording during the stimulation night. While asleep, tones associated with one of the sequences were replayed to the participants during stable N2 and N3. During S4 (20 days post-TMR) SRTT was delivered outside the scanner. (**b**) MRI data analysis. MD and Fr maps were extracted from the DW-MRI data and combined in a multimodal approach to uncover common trends related to microstructural plasticity of grey matter. Multimodal analysis was followed by unimodal post-hoc tests to determine the contribution of each modality to the multimodal results (not shown). SRTT: Serial Reaction Time Task; DW-MRI: Diffusion Weighted MRI; MD: Mean Diffusivity; Fr: Restricted Water Fraction.

Session 2 (S2), session 3 (S3) and session 4 (S4) took place 23-26 h, 10-14 days, and 16-21 days after S1, respectively. During S2 and S3 DW-MRI data were acquired as before, followed by an SRTT re-test. Here, the first half of the SRTT blocks (24 sequence blocks + 4 random blocks) was performed in the 3T scanner and the second half (24 sequence blocks + 4 random blocks) in the 0T scanner. Note that the order of scans (3T vs 0T) was flipped from S1 to S2 and S3 for the microstructural assessment to occur as close to TMR as possible. S4 was performed either in the lab or online, depending on the severity of COVID-19 restrictions at the time. During S4, SRTT was delivered in one run (48 sequence blocks + 4 random blocks).

For offline data collection, the SRTT (S1-S3) was back projected onto a projection screen situated at the end of the MRI/mock scanner and reflected into the participant’s eyes via a mirror mounted on the head coil; during S4 SRTT was presented on a computer screen with resolution 1920 × 1080 pixels and executed using MATLAB 2016b (The MathWorks Inc., Natick, MA, USA) and Cogent 2000 (developed by the Cogent 2000 team at the Functional Imaging Laboratory and the Institute for Cognitive Neuroscience, University College, London, UK; http://www.vislab.ucl.ac.uk/cogent.php). For online data collection (S4), SRTT was programmed in Python using PsychoPy 3.2.2. ^69^ and administered through Pavlovia online platform (https://pavlovia.org/).

### 5.3 EXPERIMENTAL TASKS - THE SERIAL REACTION TIME TASK (SRTT)

The SRTT was used to induce and measure motor sequence learning. It was adapted from ^17^ and implemented exactly as described in ^24^. Briefly, participants learned two 12-item sequences of auditorily and visually cued key presses. The task was to respond to the stimuli as quickly and accurately as possible, using index and middle fingers of both hands. The two sequences - A (1-2-1-4-2-3-4-1-3-2-4-3) and B (2-4-3-2-3-1-4-2-3-1-4-1) - were matched for learning difficulty, did not share strings of more than four items and contained items that were equally represented (three repetitions of each). Each sequence was paired with a set of 200 ms-long tones, either high (5^th^ octave, A/B/C#/D) or low (4^th^ octave, C/D/E/F) pitched, that were counterbalanced across sequences and participants. For each item/trial, the tone was played with simultaneous presentation of a visual cue in one of the four corners of the screen. Visual cues consisted of neutral faces and objects, appearing in the same location regardless of the sequences (1 - top left corner = male face, 2 - bottom left corner = lamp, 3 - top right corner = female face, 4 - bottom right corner = water tap). Participants were told that the nature of the stimuli (faces/objects) was not relevant for the study. Their task was to press the key on the keyboard (while in the sleep lab or at home) or on an MRI-compatible button pad (2-Hand system, NAtA technologies, Coquitlam, Canada) (while in the MRI/mock scanner) that corresponded to the position of the picture as quickly and accurately as possible: 1 = left shift/left middle finger button; 2 = left Ctrl/left index finger button; 3 = up arrow/right middle finger button; 4 = down arrow/right index finger button. Participants were instructed to use both hands and always keep the same fingers on the appropriate response keys. The visual cue disappeared from the screen only after the correct key was pressed, followed by a 300 ms interval before the next trial. There were 24 blocks of each sequence (a total of 48 sequence blocks per session), where block type was indicated with ‘A’ or ‘B’ displayed in the centre of the screen. Each block contained three sequence repetitions (36 items) and was followed by a 15s pause/break, with reaction time (RT) and error rate feedback. Blocks were interleaved pseudo-randomly with no more than two blocks of the same sequence in a row. Participants were aware that there were two sequences but were not asked to learn them explicitly. Block order and sequence replayed were counterbalanced across participants.

During each run of the SRTT, sequence blocks A and B were followed by 4 random blocks, except for the first half of S1 (to avoid interrupted learning). Random blocks were indicated with ‘R’ appearing centrally on the screen and contained pseudo-randomised sequences, the same visual stimuli, and tones matching sequence A for half of them (Rand_A) and sequence B for the other half (Rand_B). Blocks Rand_A and Rand_B were interleaved, and the random sequences contained within them followed three constraints: (1) each cue was represented equally within a string of 12 items, (2) two consecutive trials could not contain the same cue, (3) random sequence did not share a string of more than four items with either sequence A or B.

### 5.4 EEG DATA ACQUISITION

EEG was recorded using 64 actiCap slim active electrodes (Brain Products GmbH, Gilching, Germany), with 62 electrodes embedded within an elastic cap (Easycap GmbH, Herrsching, Germany). This included the reference positioned at CPz and ground at AFz. The remaining electrodes were the left and right electrooculography (EOG) electrodes (placed below and above each eye, respectively), and left and right electromyography (EMG) electrodes (placed on the chin). Fig.S2 shows the EEG electrodes layout. Elefix EEG-electrode paste (Nihon Kohden, Tokyo, Japan) was used for stable electrode attachment and Super-Visc high viscosity electrolyte gel (Easycap GmbH) was inserted into each electrode to reduce impedance below 25 kOhm. To amplify the signal, we used either two BrainAmp MR plus EEG amplifiers or a LiveAmp wireless amplifier (all from Brain Products GmbH). Signals were recorded using BrainVision Recorder software (Brain Products GmbH).

### 5.5 TMR DURING NREM SLEEP

Tones associated with one of the learned sequences (A or B, counterbalanced across participants) were replayed to the participants during N2 and N3, as assessed with standard AASM criteria ^70^. Volume was adjusted for each participant to make sure that the sounds did not wake them up. One repetition of a sequence was followed by a 20 s break, with the inter-trial interval jittered between 2500 and 3500 ms. Upon arousal or leaving the relevant sleep stage, replay was paused immediately and resumed only when stable N2/N3 was apparent. TMR was performed for as long as a minimum threshold of ∼1000 trials in N3 was reached. On average, 1552.91 ± 215.00 sounds were delivered. The protocol was executed using MATLAB 2016b and Cogent 2000.

### 5.6 MRI DATA ACQUISITION

Magnetic resonance imaging (MRI) was performed at Cardiff University Brain Imaging Centre (CUBRIC) with a 3T Siemens Connectom scanner (maximum gradient strength 300 mT/m). All scans were acquired with a 32-channel head-coil and lasted ∼1 h in total each, with whole-brain coverage including cerebellum. This paper is concerned with the analysis of the multi-shell DW-MRI, but the MRI protocol also included T1w, functional MRI (fMRI) and mcDESPOT acquisitions, the analyses of which are reported in separate publications (for T1w and fMRI results see ^24^).

#### 5.6.1 T1-WEIGHTED IMAGING

A high resolution T1-weighted anatomical scan was acquired with a 3D magnetization-prepared rapid gradient echoes (MPRAGE) sequence (repetition time [TR] = 2300 ms; echo time [TE] = 2 ms; inversion time [TI] = 857 ms; flip angle [FA] = 9°; bandwidth 230 Hz/Pixel; 256 mm field-of-view [FOV], 256 × 256 voxel matrix size, 1 mm isotropic voxel size; 1 mm slice thickness; 192 sagittal slices; parallel acquisition technique [PAT] with in-plane acceleration factor 2 (GRAPPA); anterior-to-posterior phase-encoding direction; 5 min total acquisition time [AT]) at the beginning of each scanning session.

#### 5.6.2 MULTI-SHELL DIFFUSION-WEIGHTED IMAGING

Diffusion-Weighted MRI data was acquired with a monopolar sequence (TR = 3000 ms; TE = 59 ms; FA = 90°; 266 gradient directions distributed over 6 shells (b = 200, 500, 1200, 2400, 4000, 6000 s/mm^2^); 13 interspersed b = 0 images; bandwidth 2272 Hz/Pixel; 220 mm FOV; 220 × 200 voxel matrix size; 2 mm isotropic voxel size; 2 mm slice thickness; 66 axial-to-coronal slices obtained parallel to the AC-PC line with interleaved slice acquisition; PAT 2 (GRAPPA); multi-band acceleration factor = 2; AT = 14 min) in an anterior-to-posterior phase-encoding direction, with one additional b = 0 posterior-to-anterior volume.

### 5.7 DATA ANALYSIS

#### 5.7.1 BEHAVIOURAL DATA

PSQI global scores were determined in accordance with the original scoring system ^68^ and the SRTT analysis was performed as before ^24^. Briefly, the SRTT performance was measured using mean reaction time per block of each sequence (cued and uncued). All trials within each block were considered (i.e., trials performed by both hands), except for those with reaction time exceeding 1,000 ms. For each sequence during each session, the mean performance on the last 4 blocks was subtracted from the mean performance on the 2 random blocks, thus yielding a measure of late ‘sequence-specific skill’ (SSS). We chose to focus on late SSS rather than early SSS given our prior results showing a main effect of TMR on the former only ^19,24^.

To obtain a single measure reflecting the effect of TMR on SRTT performance at each session we calculated the difference between the late SSS of the cued and uncued sequence and refer to it as the ‘cueing benefit’. Cueing benefit at S2, S3, and S4 were entered as the main regressors in the analysis testing the relationship between microstructural plasticity and cueing benefit at different sessions (see section *2.7.4.2 DW-MRI: multimodal analysis of TMR-related plasticity)*.

Finally, to obtain a single measure reflecting an overall susceptibility to the manipulation, we performed Principal Component Analysis (PCA) on the cueing benefit at S2, S3 and S4 (i.e., the three post-TMR sessions). This yielded a PCA-transformed cueing benefit, derived from the first PCA component and referred to as the ‘TMR susceptibility’. TMR susceptibility was entered as the main regressor in the analysis testing the relationship between inter-individual variability in brain microstructure and susceptibility to the manipulation (see section *2.7.4.4 DW-MRI: Individual differences in baseline (micro)structure)*.

#### 5.7.2 DW-MRI DATA PRE-PROCESSING

DW-MRI data pre-processing was performed as described in previous publications ^71,72^. The pre-processing steps included (1) Slicewise OutLIer Detection (SOLID) ^73^; (2) full Fourier Gibbs ringing correction ^74^ using Mrtrix mrdegibbs software ^75^; and (3) a combined topup, eddy and DISCO step ^76^ to (i) estimate susceptibility-induced off-resonance field and correct for the resulting distortions using images with reversed phase-encoding directions, (ii) correct for eddy current distortions and (iii) correct for gradient nonlinearity. To generate Mean Diffusivity (MD) maps, the diffusion tensor model was fitted to the data using the DTIFIT command in FSL for shells with b < 1500 s/mm^2^. To estimate Restricted Water Fraction (Fr) metric, the composite hindered and restricted model of diffusion (CHARMED) was fitted to the data using an in-house non-linear least square fitting algorithm ^77^ coded in MATLAB 2015a. The two indices (MD, Fr) were chosen based on the existing human literature on the microstructural changes following learning (MD: ^26,27,29,30^ Fr: ^28^). MD describes the average mobility of water molecules and has shown sensitivity to changes in grey matter ^26,29,30^. MD is thought to reflect the underlying, learning-dependent remodelling of neurons and glia, i.e., synaptogenesis, astrocytes activation and brain-derived neurotrophic factor (BDNF) expression, as confirmed by histological findings ^26^, which were of particular interest in this study. As opposed to DTI, the CHARMED model separates the contribution of water diffusion from the extra-axonal (hindered) and intra-axonal (restricted) space ^32^, thereby providing a more sensitive method to look at the microstructural changes than DTI ^28^. Fr is one of the outputs from the CHARMED framework. In grey matter, Fr changes are thought to reflect remodelling of dendrites and glia, and were observed both short-term (2 h) and long-term (1 week) following a spatial navigation task ^28^.

Co-registration, spatial normalisation and smoothing of the MD and Fr maps were performed in SPM12, running under MATLAB 2015a. First, we co-registered the pre-processed diffusion images with participants’ structural images using a rigid body model. The co-registration output was then spatially normalised to MNI space. This step involved resampling to 2 mm voxel with B-spline interpolation and utilised T1 deformation fields generated during fMRI analysis of the same participants ^24^. That way, the resulting diffusion images were in the same space as the fMRI and T1w data. Finally, the normalised data was smoothed with an 8 mm FWHM Gaussian kernel.

#### 5.7.3 STATISTICAL ANALYSIS

All behavioural tests conducted were two-tailed, and both positive and negative contrasts were performed for the MRI analyses. MRI results were voxel-level corrected for multiple comparisons by family wise error (FWE) correction for the whole-brain grey matter (GM) and for the pre-defined anatomical regions of interest (ROI, see section *2.7.4.6*), with the significance threshold set at p_FWE_ < 0.05. To obtain a whole-brain GM mask, the SPM12 tissue probability map of GM was thresholded at 50% probability ^78^.

##### 5.7.3.1 BEHAVIOURAL DATA

Statistical analyses of behavioural data were conducted in a prior write-up ^19^. We used lme4 package ^79^ in R to fit two linear mixed effects models to our data. The first model (model 1) was used to test the effect of TMR (cued vs uncued) and Session (S2, S3, S4) on the late SSS. The second model (model 2) was used to test the effect of Time (i.e., number of days post-TMR) on cueing benefit. To account for the repeated measures design, participant code was always entered as a random intercept.

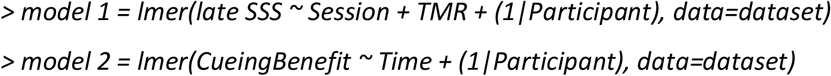

An ANOVA function in R was used to run likelihood ratio tests between the full model and the model without the effect of interest. This allowed us to obtain a p-value for each effect tested. Emmeans package ^80^ was used to conduct Holm-adjusted post-hoc pairwise comparisons. The results of both the likelihood ratio tests and post-hoc comparisons are cited in this manuscript and discussed in relation to the underlying tissue microstructure.

##### 5.7.3.2 MULTIMODAL ANALYSIS

Group level analyses of DW-MRI data was performed in FSL (FMRIB’s Software Library, http://www.fmrib.ox.ac.uk/fsl) ^81^. To examine the relationship between brain characteristics and our variables of interest we performed non-parametric combination (NPC) for joint interference analysis (Fig.5b), as described before ^82,83^. Specifically, NPC was performed over MD and Fr maps to uncover common trends related to non-myelin GM microstructure ^26,28^.

The analysis was performed through Permutation Analysis of Linear Models (PALM) in FSL ^84^, using voxel-wise Fisher test with the following equation:

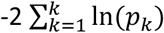

where k denotes the total number of modalities being combined, and p_k_ denotes the p-value for a given modality ^84^.

NPC Fisher’s combining function tests for effects with concordant directions across modalities of choice. Thus, to test for positive effects across our modalities, imaging data with mismatching directions (here MD) were multiplied by (−1). The significance of the resulting, single joint statistic was assessed through 5000 permutations of each of the separate tests and a cluster-forming threshold of t > 1.75 (equivalent to p < 0.05, based on the degrees of freedom for the smallest sample) at 5% FWE rate. Correction for multiple comparisons was carried out both for the whole brain GM and for the pre-defined ROIs.

Having previously established that TMR can impact on GM volume ^24^, here we investigated the relationship between early (S1 vs S2) and late (S2 vs S3) microstructural plasticity and cueing benefit at different time points. To this end, the images used to assess the longitudinal effects of time were generated by subtracting the pre-processed MD and Fr parameter maps from consecutive sessions. The resultant images were entered into a one-sample t-test with cueing benefit at S2, S3 and S4 added as regressors. We run separate contrasts for each session, whereby the session of interest was specified as the main regressor whereas the remaining sessions were treated as the covariates of no interest (nuisance covariates). This ensured that the results were specific to the session analysed. Additionally, the nuisance covariates also included sex and age to control for the differences between males and females, as well as the effect of age. Baseline reaction time (i.e., average reaction time on the random blocks performed during S1) and baseline learning capabilities (i.e., difference between the average of the last 4 blocks and the first 4 blocks performed during S1) were also specified as the variables of no interest to ensure that the results were independent of baseline SRTT performance.

To determine whether individual differences in baseline brain characteristics can predict susceptibility to the manipulation we tested the relationship between baseline (S1) GM microstructure and the variance shared between cueing benefit at the post-TMR sessions. Thus, the PCA-transformed cueing benefit, here referred to as the ‘TMR susceptibility’ was entered as a covariate of interest in a one-sample t-test within the multimodal framework. The nuisance covariates for this analysis included: sex, age, PSQI, baseline reaction time, baseline learning capabilities, and percentage of time spent in N2 (given the results described in ^85^). This approach ensured that the results were independent of demographics, general sleep patterns and baseline SRTT performance, all of which could be related to baseline characteristics of the brain.

##### 5.7.3.3 UNIMODAL ANALYSIS

To determine individual contribution of each microstructural modality to the multimodal results, we performed unimodal analyses of individual modalities, in FSL and through non-parametric, permutation-based voxel-wise comparisons using the *randomise* function ^86^. Results were derived from 5000 permutations. Correction for multiple comparisons was carried out by FWE correction both for the whole brain GM and for the pre-defined ROIs, as for the multimodal analysis. Multiple modalities correction was performed based on the number of modalities entered in the multimodal analysis (here two: MD and Fr).

##### 5.7.3.4 REGIONS OF INTEREST

All MRI results were voxel-level corrected for multiple comparisons within the whole-brain GM and the pre-defined anatomical ROIs. Our pre-defined ROIs included (1) bilateral precuneus, (2) bilateral hippocampus and parahippocampus, (3) bilateral dorsal striatum (putamen and caudate), (4) bilateral cerebellum, (5) bilateral sensorimotor cortex (precentral and postcentral gyri). All ROIs were selected based on their known involvement in sleep-dependent procedural memory consolidation ^87-90^ and memory reactivation ^13,18,22,30,91,92^). A mask for each ROI was created using an Automated Anatomical Labeling (AAL) atlas in the Wake Forest University (WFU) PickAtlas toolbox ^93^.

### 5.8 RESULTS PRESENTATION

Anatomical localisation of the significant clusters from both unimodal and multimodal analyses was determined with the automatic labelling of MRIcroGL (https://www.nitrc.org/projects/mricrogl/) based on the AAL atlas. Results Fig.2-5 and Fig.S3-S4 are presented using MRIcroGL, displayed on the MNI152 standard brain (University of South Carolina, Columbia, SC). All significant clusters are reported in supplementary tables, but only those with an extent equal to or above 5 voxels are discussed in text and presented in figures. Fig.5, Fig.S1 and Fig.S2 were created in Microsoft PowerPoint v16.53, Fig.1a was generated using *ggplot2* (version 3.3.0) ^94^ in R, Fig.1b was generated using Prism 9 (GraphPad Software, San Diego, CA, USA), and Fig.1c was generated using *corrplot* command in MATLAB.

## DATA AND CODE AVAILABILITY

All data collected during the study, scripts that delivered experimental tasks and codes used to conduct the analyses are publicly available at: DOI 10.17605/OSF.IO/B52FV.

## ACKNOWLEDGEMENTS

The authors would like to thank Holly Kings and Sofia Pereira for helpful comments and discussion on this manuscript, Eleonora Patitucci, Sonya Foley, and Marco Bigica for advice on the MRI analysis, as well as Mahmoud E. A. Abdellahi, Chelsea Bryant, and Joe Davis for support with participants’ recruitment and data collection. The authors are also grateful to Jennifer Roebber for sharing her SRTT script on Pavlovia and her help with Python coding. We further acknowledge Chantal Tax and Greg Parker who developed the Modular Pipeline for Multishell DTI processing used in this study. Finally, we thank all the participants for their time and commitment to the study. This work was supported by the ERC grant SolutionSleep, 681607, to PL.

## AUTHORS CONTRIBUTION STATEMENT

M.R. and P.A.L. conceived the study and designed the experiment, M.R., and P.B. acquired the MRI data; M.R., A.L., M.C. and P.B. performed the analysis; M.R wrote the manuscript with input from all co-authors; P.A.L. supervised the project and obtained funding.

