## Supplementary Materials for "Distributed and gradual microstructure changes track the emergence of behavioural benefit from memory reactivation"

### SUPPLEMENTARY TABLES:

***Table S1. Microstructural plasticity associated with cueing benefit at different sessions.***

*Cluster statistics for precuneus, putamen and sensorimotor cortex which showed a positive relationship microstructural plasticity (MD and Fr) and cueing benefit at different sessions. MD and Fr were either analysed together in a multimodal framework (i) or separately using unimodal analyses (ii, iii).*

| Analysis | ROI | Region | MNI x, y, z (mm) | Number of voxels | T peak | $P_{FWE}$ peak |
|---|---|---|---|---|---|---|
| **A.** [Early microstructural plasticity * cueing benefit at S3] | | | | | | |
| i) Multimodal | Precuneus | Left precuneus | -6, -60, 18 | 1328 | - | **0.041*** |
| ii) Unimodal: MD | | - | -10, -58, 30 | 62 | 2.65 | 0.619 |
| iii) Unimodal: Fr | | Left precuneus | -2, -56, 8 | 1378 | 8.02 | 0.077 |
| **B.** [Late microstructural plasticity * cueing benefit at S4] | | | | | | |
| i) Multimodal | Precuneus | Right precuneus | 4, -58, 16 | 1943 | - | **0.027*** |
| ii) Unimodal: MD | | Right precuneus | -6, -54, 30 | 1992 | 10.08 | **0.033*** |
| iii) Unimodal: Fr | | - | 8, -52, 6 | 867 | 8.93 | 0.116 |
| **C.** [Early microstructural plasticity * cueing benefit at S4] | | | | | | |
| i) Multimodal | Dorsal Striatum | Right putamen | 34, -6, -8 | 633 | - | **0.016*** |
| ii) Unimodal: MD | | Right putamen | 32, 2, -10 | 563 | 5.20 | 0.054^ |
| iii) Unimodal: Fr | | - | 20, 8, -4 | 79 | 3.34 | 0.563 |
| **D.** [Late microstructural plasticity * cueing benefit at S4] | | | | | | |
| i) Multimodal | Sensorimotor Cortex | Left precentral and postcentral gyri | -60, -18, 14 | 2159 | - | **0.018*** |
| ii) Unimodal: MD | | Left precentral and postcentral gyri | -60, -20, 14 | 3987 | 13.15 | **0.010*** |
| iii) Unimodal: Fr | | - | 48, -8, 56 | 54 | 7.71 | 0.783 |

*Regions listed were significant at peak voxel threshold of $p_{FWE} < 0.05$, after correction for multiple voxel-wise comparisons within pre-defined bilateral ROI (as listed in the first column) and the number of modalities (two modalities, MD and Fr). Peak voxel MNI coordinates and peak T values are given. Covariates of no interest included in the analysis: age, sex, PSQI score, baseline reaction time, baseline learning capabilities on the SRTT, cueing benefit at S2, cueing benefit at S4 (for A only), cueing benefit at S3 (for B-D). S3-4: Session 3-4; * $p < 0.05$, ^ $p < 0.06$. n = 16 for (A, C), n = 15 for (B, D).*

***Table S2. Baseline microstructure and TMR susceptibility.***

*Cluster statistics for sensorimotor cortex which showed a positive relationship between baseline microstructure and TMR susceptibility (i.e., PCA-transformed cueing benefit at session 2, 3 and 4).*

| Analysis | ROI | Region | MNI x, y, z (mm) | Number of voxels | T peak | $P_{FWE}$ peak |
|---|---|---|---|---|---|---|
| i) Multimodal | Sensorimotor Cortex | Left precentral and postcentral gyri | 58, -6, 20 | 1691 | - | **0.041*** |
| ii) Unimodal: MD | | - | 54, 0, 18 | 1396 | 9.42 | 0.052^ |
| iii) Unimodal: Fr | | - | -34, -28, 40 | 233 | 5.00 | 0.219 |

*Regions listed were significant at peak voxel threshold of $p_{FWE} < 0.05$, after correction for multiple voxel-wise comparisons within pre-defined bilateral ROI (as listed in the first column) and the number of modalities (two modalities, MD and Fr). Peak voxel MNI coordinates and peak T values are given. Covariates of no interest included in the analysis: age, sex, PSQI score, baseline reaction time and baseline learning capabilities on the SRTT, percentage of time spent in N2. * $p < 0.05$, ^ $p < 0.06$. n = 16.*

### SUPPLEMENTARY FIGURES:

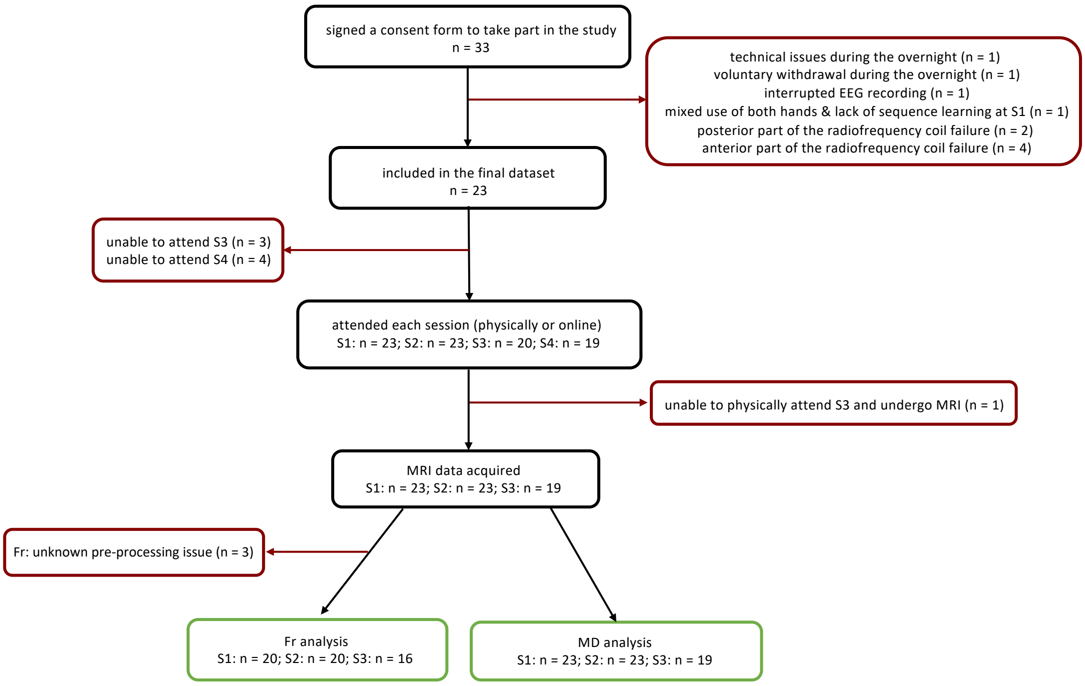


***Fig. S1. A flowchart showing participants included and excluded from the analysis.*** *In black and white, the number of participants included in the study at different time points, with the final sample size shown in green. In red, the number of participants excluded from different analyses, together with a reason for the exclusion. S1-S4: Session 1 – Session 4; Fr: Restricted water fraction; MD: Mean Diffusivity.*

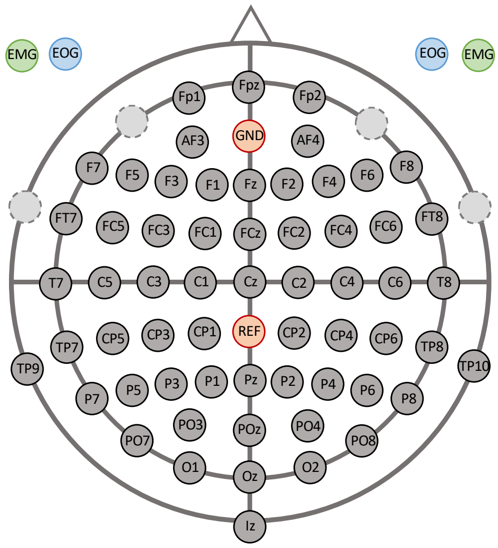


***Fig. S2. EEG electrodes layout****. Orange: ground (GND) and reference (REF) electrodes; light grey (dashed circles): original position of the electrodes used to record EMG and EOG; green: EMG electrodes; blue: EOG electrodes.*

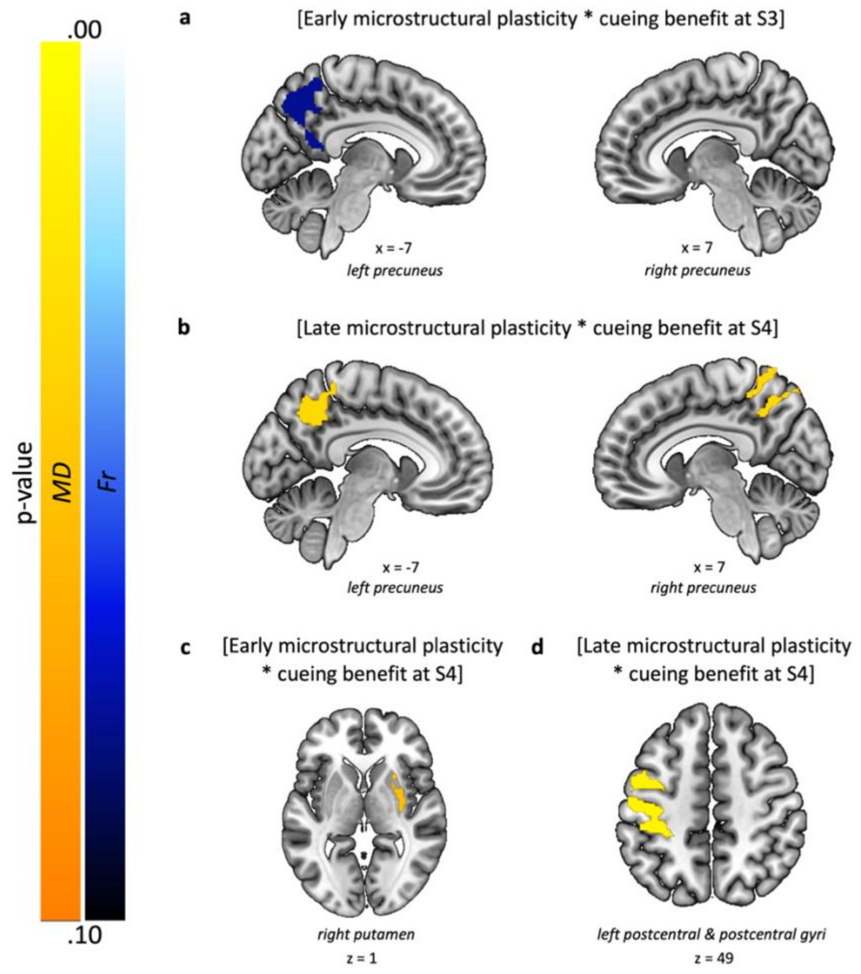


***Fig. S3. Results of unimodal analyses conducted for MD and Fr separately, testing the same relationships as in Fig.2-4***. *(a, b) Unimodal results for the same contrasts as in Fig.2a and Fig.2d, respectively. (**c, d**) Unimodal results for the same contrasts as in Fig.3a and Fig.3d, respectively. Colour bars indicate p-values at $p_{FWE} < 0.1$, corrected for multiple modalities and the number of voxels within the chosen ROI. In orange, MD clusters; in blue, Fr clusters. Results are overlaid on a Montreal Neurological Institute (MNI) brain. Colour bars indicate 1 - p-value, derived from 5000 permutations. S1-4: Session 1-4. n = 16 for (a, c), n = 15 for (b, d).*

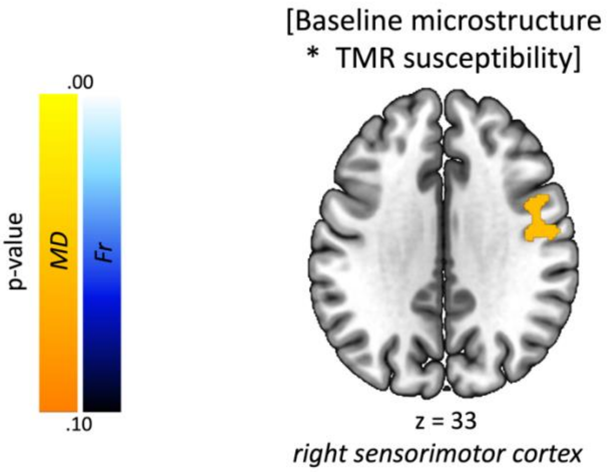


***Fig. S4. Results of unimodal analyses conducted for MD and Fr separately, testing the same contrast as in Fig.4**. Colour bars indicate p-values at $p_{FWE} < 0.1$, corrected for multiple modalities and the number of voxels within the chosen ROI. In orange, MD clusters; in blue, Fr clusters. No Fr clusters above the $p_{FWE} < 0.1$ threshold were revealed. Results are overlaid on a Montreal Neurological Institute (MNI) brain. Colour bars indicate 1 - p-value, derived from 5000 permutations. S1-4: Session 1-4. n = 16.*
